# Activity-Dependent Postsynaptic Mitochondrial ROS Signaling Drives Avoidance Plasticity *in C. elegans*

**DOI:** 10.64898/2026.06.05.730457

**Authors:** Kaz M. Knight, Zephyr Lenninger, Ennis Deihl, Rachel L. Doser, Frederic J. Hoerndli

## Abstract

Reactive oxygen species (ROS) are signaling molecules involved in neuronal excitatory function, with mitochondrial ROS (mitoROS) playing key roles in metabolic regulation and stress responses. Studies have shown that neuronal activity upregulates mitoROS production through oxidative phosphorylation, but it remains unclear if and how acute elevations in mitoROS influence synaptic plasticity. Here, we develop an avoidance sensitization paradigm in *C. elegans* using optogenetic excitation and training of nociceptive ASH neurons to initiate avoidance reversals by downstream activation of the AVA command interneurons. Using this paradigm, we show that the probability of reversal to light stimulation increases 4-hours after optogenetic training, indicating behavioral sensitization. This avoidance sensitization is accompanied by an increase of surface glutamate receptor (GLR-1) levels at ASH-AVA synapses which is dependent on postsynaptic expression of GLR-1 and active transcription. Interestingly, we find that somatic and nerve ring mitochondria produce ROS after optogenetic training. We show that this mitochondrial ROS (mitoROS) peak is dependent on postsynaptic GLR-1 and MCU-1 function during optogenetic training and is necessary for avoidance sensitization. Finally, we demonstrate that postsynaptic signaling by mitoROS in AVA is sufficient to induce avoidance sensitization. Postsynaptic photoactivation of mitochondria-targeted Killer Red in AVA, calibrated to produce the mitoROS peak observed during training, induces avoidance sensitization bypassing optogenetic training and MCU-1 requirement. Our results indicate that activity-dependent mitoROS signaling can instruct synaptic strengthening and directly modulate circuit function and behavior.

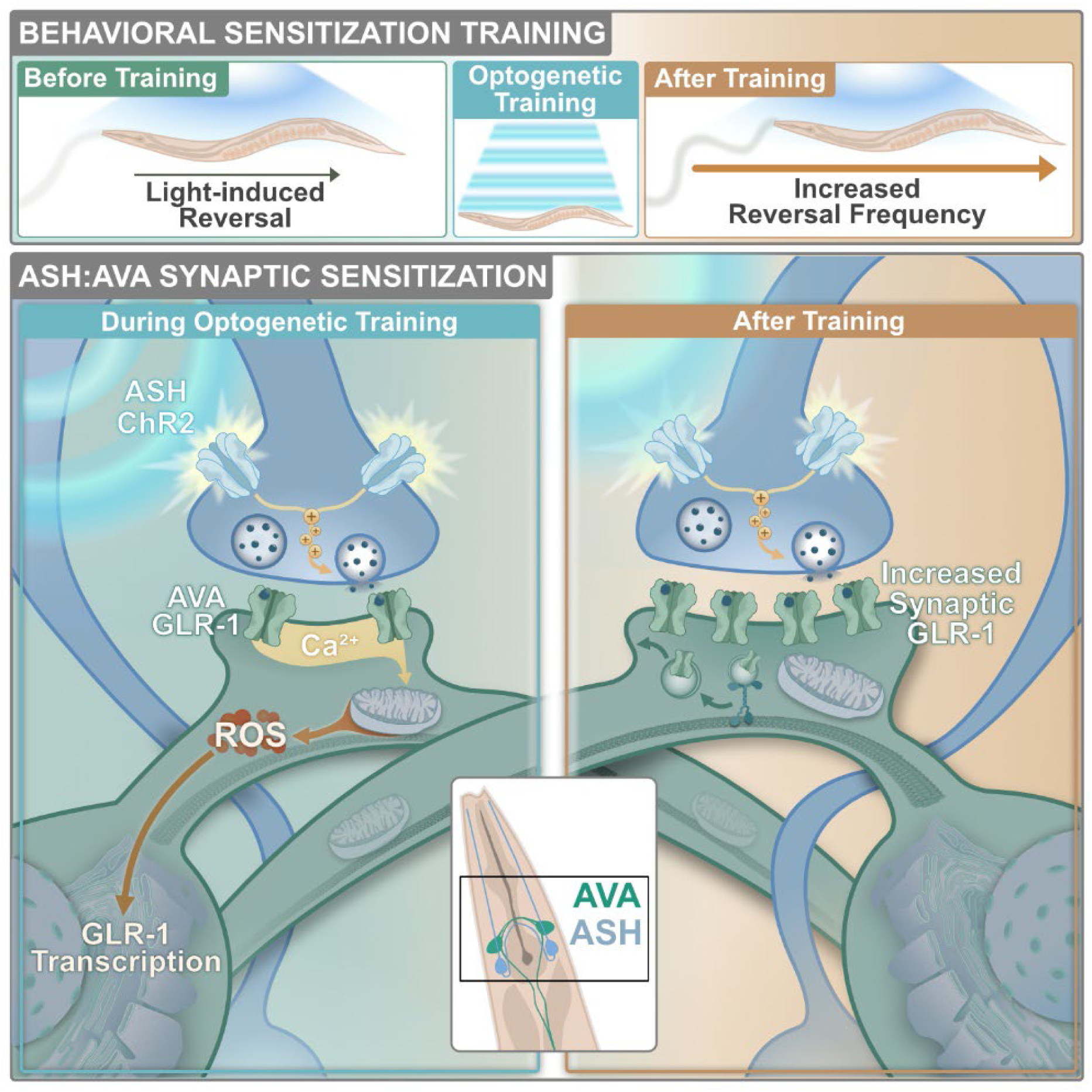

## Introduction

Neuronal function, especially synaptic transmission and plasticity, demands considerable amounts of energy.^1^ Combined with spatiotemporal complexity, neurons present unique challenges, requiring the hub-like utility of mitochondria to adapt to local and cell-wide demands.^2,3^ Proper regulation of mitochondrial position and morphology contributes to compartmentalization of the postsynaptic space and supports the dynamic demands of each compartment.^4–7^ Additionally, mitochondria have critical roles beyond energy production, including regulating calcium handling, redox signaling, and stress responses.^8–11^ There is accumulating evidence that these roles are instrumental to synaptic development,^12–14^ plasticity^1,8,15–20^ and adaptation during neurodegenerative disease.^12,14,17,21,22^ However, given their multifunctional roles, deciphering exactly how mitochondria are involved in these key synaptic functions, especially *in vivo*, is challenging.

Mitochondrial function is closely tied to calcium flow through the mitochondria calcium uniporter (MCU-1).^23–25^ In neurons, increases in activity lead to calcium influx into mitochondria through MCU-1, which buffers cytoplasmic calcium and modulates oxidative phosphorylation resulting in increased ATP and reactive oxygen species (ROS) production.^20,26^ Additionally, this upregulated mitochondrial function is known to drive signaling cascades involved in cell death, glutamate receptor expression and long-term potentiation (LTP).^27,28^ Persistent increases in activity can result in more drastic mitochondrial morphology changes through fusion/fission or mitophagy in neurodegeneration.^29–31^ However, the functional significance of activity-dependent changes in mitochondria to circuit plasticity and behavioral adaptations is not well understood. This gap is primarily due to the lack of data from *in vivo* models where the relationship between synaptic plasticity and behavioral changes can be directly assessed.

Sensitization is characterized by an increased response to a repeated stimulus. Early studies from Eric Kandel using Aplysia provided the groundwork linking behavioral sensitization to postsynaptic ionotropic receptors, calcium (Ca^2+^) signaling and gene transcription.^32,33^ Continued investigations to decipher the relationship between increased responsiveness and neuronal function have focused on the roles of pre-synaptic neurotransmitter and neuromodulator release as well as the activation of signaling cascades for enhanced neuronal responses.^34^ As stimulus training protocols are extended in intensity and duration, different long-term effects begin to surface, reliant on persistent facilitation of signaling cascades and transcription factor activation. ^35–37^ Recent addiction studies using vertebrate circuit models have also pointed to the role of postsynaptic AMPA receptor (AMPAR) expression changes in long-term sensitization.^38,39^ Finally, sensitization is often supported by simple neuronal circuits, and has been extensively characterized and studied across different model organisms with many signaling mechanisms having been found to be conserved across models.^40–42^ Taken together, sensitization and underlying synaptic signaling mechanisms provide a robust framework to understand if and how mitochondria participate in synaptic plasticity.

*Caenorhabditis elegans* is a popular *in vivo* model for investigating the role of mitochondria in neuronal function,^43,44^ neurodegeneration,^45,46^ and aging.^47–49^ The impact of mitochondrial functional adaptations in *C. elegans* and their effects on behavior mirror results seen in vertebrate models cited above.^13^ *C. elegans* has been used extensively to identify gene products and synaptic mechanisms related to well defined behaviors.^50–52^ The *C. elegans* synaptic connectome provides a unique advantage; precise mapping of synaptic inputs onto postsynaptic locations enables detailed subcellular characterization of both mitochondria and postsynaptic receptors.^53,54^ Avoidance reversal behavior has been most extensively studied because of simple and reproducible behavioral outputs (reversal length or reversal probability).

These defined outputs depend on activity of presynaptic ASH sensory neurons and downstream signaling to AVA command interneurons.^51,55,56^ Importantly, the proteins and signaling pathways involved in these mechanisms are highly conserved. ^57–64^ Finally, the transparency of *C. elegans* provides an ideal platform to investigate the mechanisms of behavioral training paradigms using optogenetic tools and fluorescent sensors and synaptic markers. We therefore decided to use *C. elegans* as an intact *in vivo* model to investigate the role of activity-dependent mitochondria function and adaptation during sensitization.

In this study, we establish a training protocol for behavior sensitization using optogenetic activation of the ASH and AVA synaptic connection important for C. *elegans* avoidance behavior (reversal of the animal). To understand the role of mitochondrial function in sensitization and synaptic plasticity, we quantified synaptic GLR-1 receptors (the *C. elegans* homolog of mammalian AMPARs), mitochondria morphology and mitoROS production in AVA command interneurons in wildtype animals. Measurements were taken before, during and after training at different subcellular locations, and compared to genetic mutants with reduced synaptic activity or mitochondrial function, as well as with cell-specific rescues. Using this approach, we quantitatively associate behavioral sensitization with mitochondrial and synaptic changes. We found that postsynaptic *mcu-1* is required for reversal sensitization and that activity-driven mitochondria adaptations depend on proximity to active synapses throughout the AVA neurite. Then, to untangle calcium signaling from mitochondrial ROS signaling, we used optogenetic activation of KillerRed (KR), a ROS producing photosensitizer, localized to mitochondria in AVA (mitoKR).^65^ We combined sensitization testing and optogenetic activation of mitoKR to show that mitoROS is necessary and sufficient for avoidance sensitization. This study demonstrates a link between neuronal activity, mitoROS production, synaptic plasticity and behavioral outcomes. As opposed to observed negative signaling roles of ROS accumulation in disease development and aging,^66^ this data points towards an instructive signaling role for ROS that strengthens active postsynaptic zones during sensitization.

## Results

### GLR-1 expression in AVA interneurons is necessary for reversal sensitization

To investigate postsynaptic mitochondrial change during persistent neuronal activity in an intact *in vivo* system, we turned to *C. elegans* avoidance and reversal behavior.^51,56^ Initiation of this behavior predominantly relies on the ASH neurons, a pair of nociceptor sensory neurons that activate during chemical, osmotic, and mechanical stimuli.^67,68^ The ASH neurons transmit the signal to motor neurons via direct and indirect connections with AVA command interneurons (Figure 1A).^56^ To gain spatiotemporal control of reversal activation, we used a previously established *C. elegans* reagent with channelrhodopsin 2 (ChR2),^56^ expressed in ASH in the *lite-1* loss-of-function (lf) background.^69,70^ Our optogenetic stimulation protocol consisted of a 1-second pulse of 470nm blue light at 200 µW/mm^2^ (Figures S1A-E) every 1 minute with a total of 5 stimulations. This led to a consistent reversal probability of 0.33 ± 0.044 reversals/stimulations (rev/stim) in control animals (Figure 1C) without depression of reversal probability as we observed for longer stimulations (2-3 seconds, Figure S1E).

**Figure 1:**
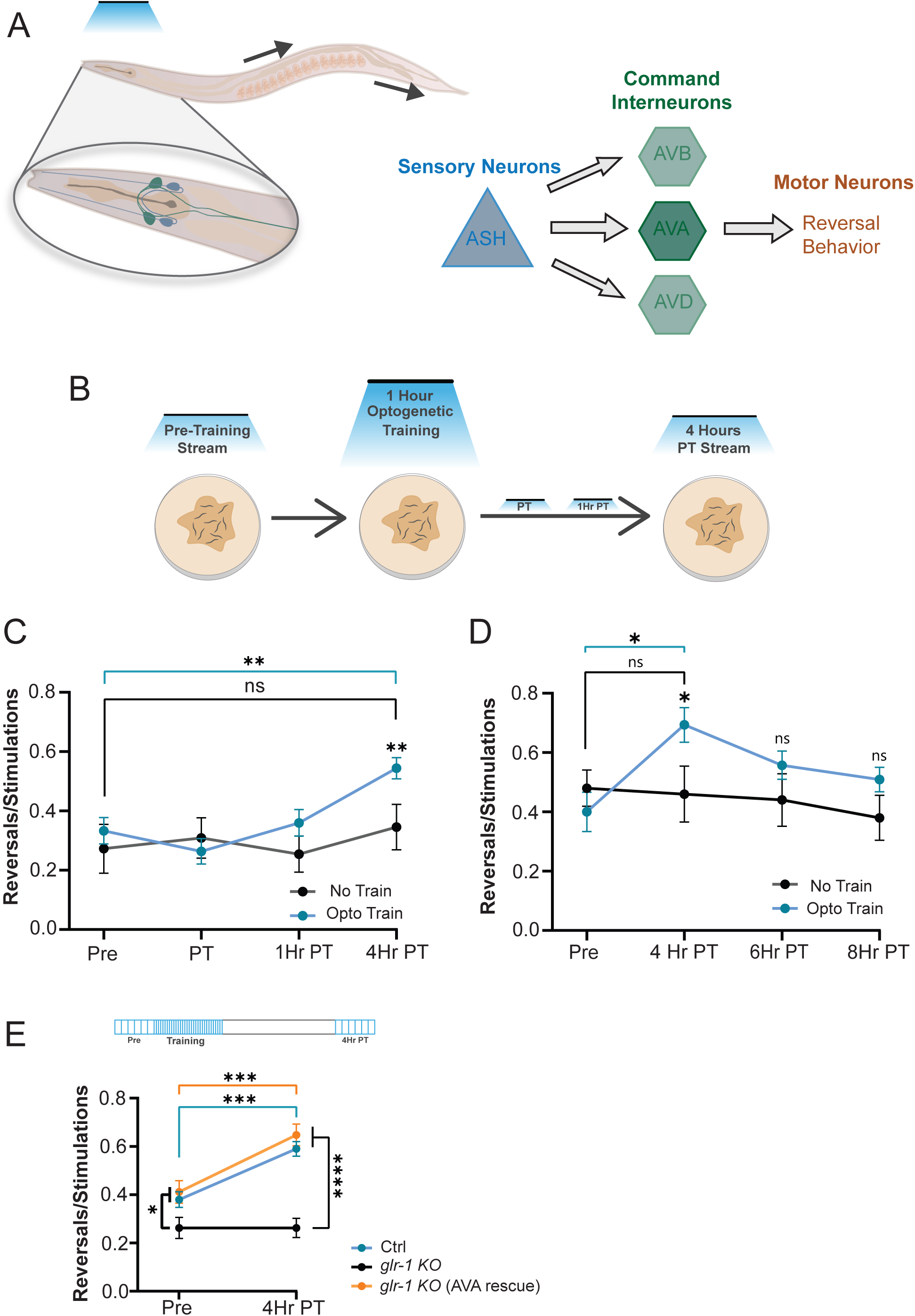
Reversal sensitization is mediated by ASH-AVA glutamatergic transmission. (A) Anatomical illustration of ASH and AVA neurons (left) and circuit diagram (right) of neurons relevant to reversals. (B) Optogenetic training and stream capture protocol for reversal sensitization. (C) Reversals/optogenetic stimulations ratio in untrained controls (n = 11) and trained animals (n ≥ 24) for pre-training, immediately post-training, 1-hour post-training, and 4-hours post-training time points. (D) Reversals/optogenetic stimulations ratio in trained animals for pre-training, 4-hours post-training, 6-hours post-training, and 8-hours post-training time (n ≥ 10). (E) Reversals/optogenetic stimulations ratio in control (n ≥ 15 animals), *glr-1(KO)* (n = 16), and *glr-1(KO)* (AVA Rescue) (n ≥ 17), animals for pre-training and 4-hours post-training time points. Data is represented as mean ± sem; ns: not significant, *p<0.05, **p<0.005, ***p<0.0005 compared to controls using a two-way ANOVA with Tukey’s correction.

To induce sensitization of the reversal circuit, we used a 1-hour training protocol consisting of a 1-second pulse of blue light every minute for an hour (Figure 1B). Videos were initially taken pre-training, post-training (PT), 1-hour post-training (1Hr PT), and 4-hours post-training (4Hr PT; Figures S1C and S1E) to detect changes in reversal counts (Figure 1C). After training, reversal ratios increased from 0.333 ± 0.044 (pre-training) to 0.552 ± 0.037 (4Hr PT). To assess longevity of the sensitization, we repeated the experiment and recorded behavior out to 8 hours post-training and found that the increase in reversal probability faded after 4-hours to 0.509 ± 0.041 8Hr PT (Figure 1D).

GLR-1 expression in AVA is necessary for spontaneous reversal behavior in *C. elegans*^60^ and learning and memory.^71–74^ In the nerve ring, there are glutamatergic ASH-AVA synapses.^75^ To test the requirement of GLR-1 for reversal sensitization, we performed optogenetic training on a global *glr-1* knockout (KO) in *lite-1(lf)* background with ASH ChR2 as described above. Untrained *glr-1(KO)* animals showed a significantly lower reversal probability to blue light before training (0.262 rev/stim ± 0.043) and did not exhibit sensitization at 4-hours PT, (0.262 ± 0.039; Figure 1E). To test cell specificity, we rescued *glr-1(KO)* in the AVA interneurons with a single copy of fluorescent-tagged GLR-1, and AVA GLR-1 expression was sufficient to restore reversal behavior sensitization (0.411 rev/stim ± 0.046 Pre to 0.648 ± 0.045 4Hr PT) (Figure 1E) indicating that ASH-AVA sensitization depends on postsynaptic expression of GLR-1 in AVA.

### Transcription-dependent increases in GLR-1 at ASH-AVA synapses are necessary for reversal sensitization

Previous studies have shown the need for GLR-1 in consolidation of sensory information,^71,73,74,76^ so we next investigated changes in synaptic GLR-1 content before and after optogenetic training by quantifying synaptic surface GLR-1 levels in the nerve ring region of AVA (in which ASH connections are reported)^53^ with and without ASH training. More specifically, we performed confocal *in vivo* imaging on strains expressing a single copy of an N-terminal fusion of GLR-1 with super-ecliptic pHluorin (SEP::GLR-1) in AVA (see methods and resources for details). Because SEP is pH sensitive, its fluorescence is quenched when GLR-1 is in trafficking vesicles or endosomal pools, thus SEP::GLR-1 fluorescence provides a measure of surface-localized receptors (Figure 2A). Using this genetic reagent, we found that total synaptic abundance (346.3 ± 35.38 to 585.7 ± 102.7; Figure 2B) and maximal intensity (458.9 ± 37.74 to 536.9 ± 53.21 ; Figure 2C) of SEP::GLR-1 in the nerve ring increased 4-hours after optogenetic training.

**Figure 2:**
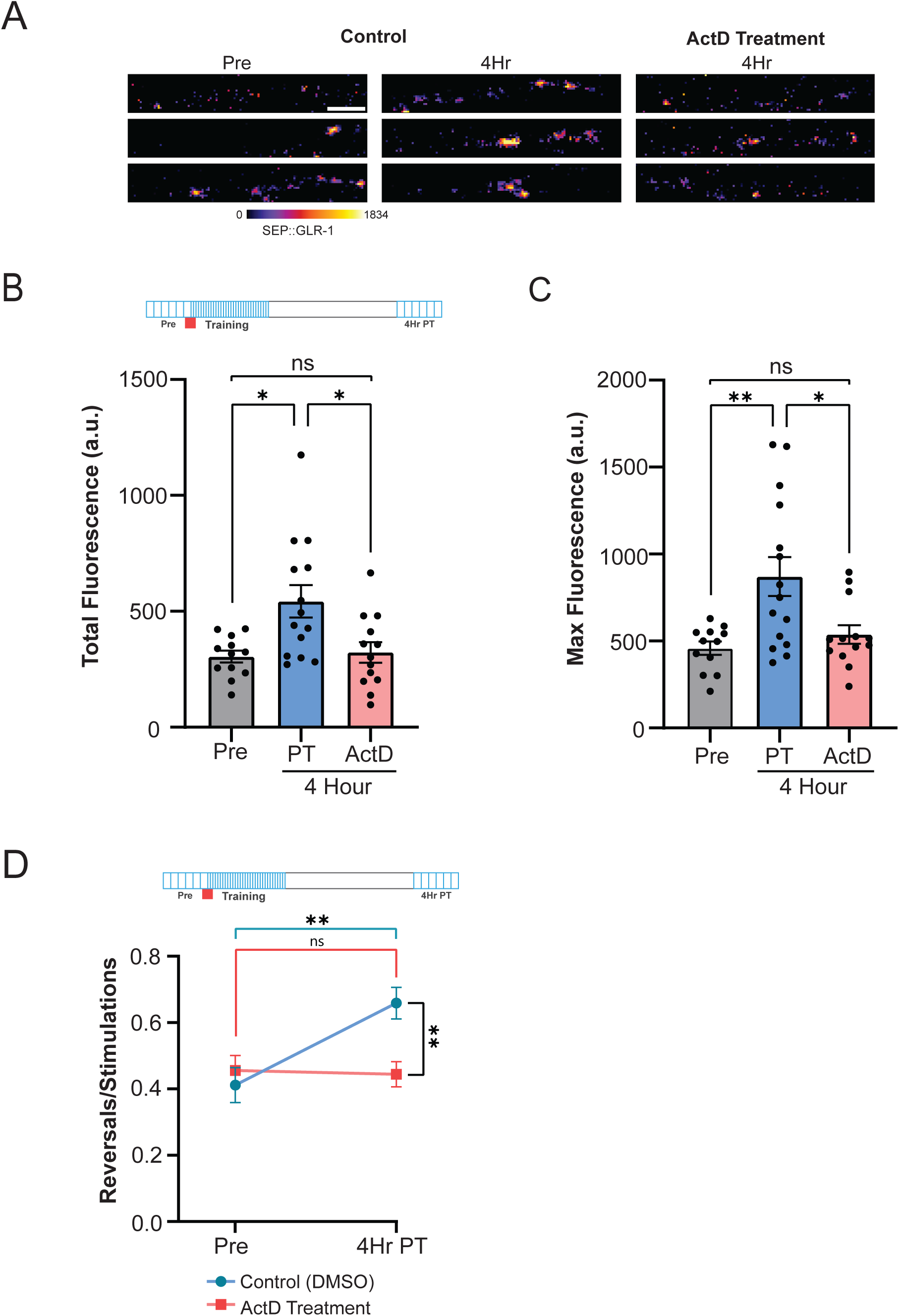
Transcription-dependent increases in GLR-1 at ASH-AVA synapses are necessary for reversal sensitization. (A) Representative images of GLR-1::SEP fluorescence in control and actinomycin D (ActD) treated animals at AVA nerve ring at pre-training and 4-hours post-training time points. Scale bar = 2 µm. (B) Total fluorescence (arbitrary units [a.u.]) or (C) maximum fluorescence of GLR-1::SEP at AVA nerve ring in control (n ≥ 12) and actinomycin D treated (n = 13) animals for pre-training and 4-hours post-training time points. Data is represented as mean ± sem; ns: not significant, *p<0.05, **p<0.005 compared to controls using a one-way ANOVA with Dunnett’s test. (D) Reversals/optogenetic stimulations ratio in control (DMSO) (n ≥ 24 animals) and actinomycin D treated (n = 18 animals) animals for pre-training and 4-hours post-training time points. Data is represented as mean ± sem; ns, not significant, **p<0.005 compared to controls using a two-way ANOVA with Tukey’s correction.

Our results indicate that the molecular changes underlying reversal sensitization occur over a 4-hour period and previous investigations into long-term sensitization indicate that behavioral changes require transcription,^77–79^ pointing toward a transcriptionally mediated mechanism. To test the hypothesis that optogenetic training and reversal sensitization requires transcription, we first verified that we could acutely block transcription with Actinomycin D (ActD) in *C. elegans.* To this end we used a reporter strain (ST66, see key resources table)^80^ that expresses GFP under the control of an heat shock inducible promoter (ncIs17[*hsp16-2p::eGFP+pBluescript]*)^80–82^ and tested whether ActD could efficiently block heat-shock induced GFP expression treatment. We saw that administering higher ActD concentrations (1000 μg/mL) could nearly immediately inhibit ∼75% of transcription of the GFP reporter (Figure S2A). With this validation, we then treated wildtype animals with ActD prior to optogenetic training (Figure 2B Top) and found that ActD treatment prevented training-dependent increases in nerve ring SEP::GLR-1 puncta (Figures 2A-C) and reversal sensitization compared to DMSO control animals (0.455 rev/stim ± 0.045 to 0.444 ± 0.038; Figure 2D). Altogether, these results suggest that synaptic strengthening and behavioral sensitization require transcriptional upregulation of GLR-1.

### Postsynaptic MCU-1-dependent mitochondrial calcium uptake is necessary for reversal sensitization and synaptic GLR-1 recruitment

Having established a robust sensitization paradigm using optogenetic stimulation of ASH, we started investigating if mitochondria participated in this form of synaptic plasticity in *C. elegans*. Our observations above suggest that glutamatergic inputs to AVA from ASH are important for our sensitization paradigm. In addition, we observed mitochondria co-localized and in close proximity to SEP::GLR-1 in the nerve ring processes of AVA where we would expect ASH-AVA synapses (Figure S3A). Our previous findings show that MCU-1 was critical for activity-dependent mitochondrial calcium uptake, production of ROS and regulation of GLR-1 trafficking.^83^ Thus, we used an *mcu-1(ju1154)* loss-of-function mutant (*mcu-1(lf)*) to eliminate activity-dependent calcium influx to the inner mitochondrial matrix without affecting basal calcium homeostasis in mitochondria.^23,83^ This mutant was indistinguishable from wildtype during pre-training, but *mcu-1(lf)* mutants did not sensitize 4-hours post training (0.3 rev/stim ± 0.045 to 0.368 ± 0.045 respectively; Figure 3A) suggesting that mitochondrial calcium uptake is necessary for increased reversals after training. To test whether calcium uptake by postsynaptic or presynaptic mitochondria is required for ASH-AVA plasticity, we performed cell-specific rescues in *mcu-1(lf)* mutants via a single-copy genomic insertion of untagged *mcu-1* expressed in either ASH or AVA.^84^ We verified rescue of mitochondrial MCU-1 function in AVA by measuring mitochondrial calcium influx using inner mitochondrial matrix-targeted GCaMP6 (Figures S3B-C).^83^ A single-copy of *mcu-1* in AVA rescued activity-dependent mitochondrial calcium uptake, so we next tested if the single-copy AVA rescue in the *mcu-1(lf)* background is sufficient for reversal sensitization. Indeed, the single-copy *mcu-1* AVA rescue fully restored reversal sensitization after 4-hours (0.388 ± 0.041 rev/stim to 0.561 ± 0.042; Figure 3A) whereas rescue in presynaptic ASH did not (Figure S3D). These results suggest that mitochondria calcium uptake by MCU-1 in postsynaptic mitochondria is necessary for reversal sensitization.

**Figure 3:**
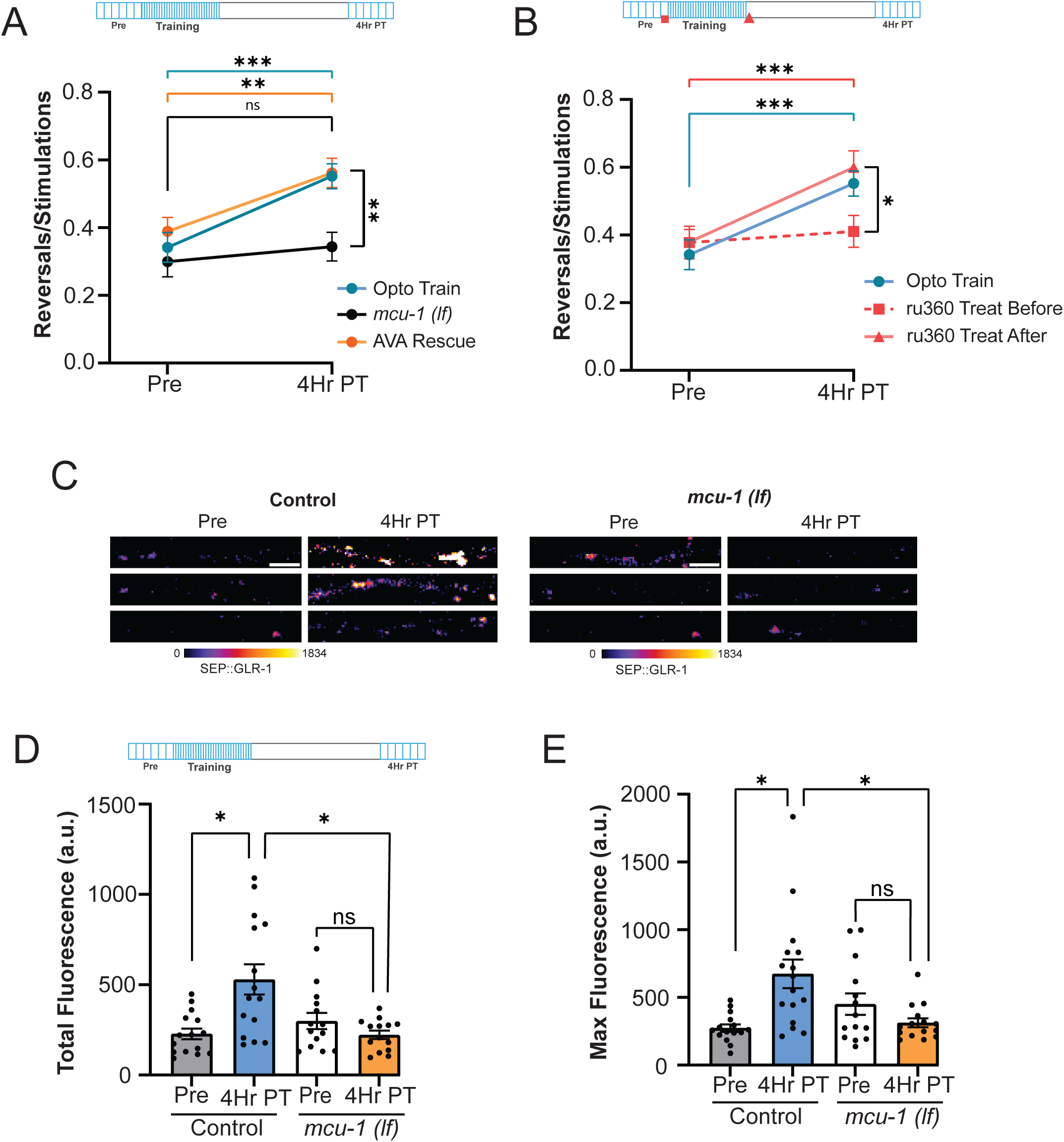
Mitochondrial calcium uptake via MCU-1 is required for training-dependent changes in GLR-1 content and reversal sensitization. (A) Reversals/optogenetic stimulations ratio in controls (n ≥ 13), *mcu-1(lf)* (n ≥ 25) and *mcu-1(lf)* (AVA Rescue) (n ≥ 22) animals at pre-training and 4-hours post-training time points. (B) Reversals/optogenetic stimulations ratio in no treatment controls (n ≥ 24 animals) and animals receiving ru360 treatment before (n ≥ 18) or after training (n ≥ 19 animals) at pre-training and 4-hours post-training time points. Data is represented as mean ± sem; ns: not significant, *p<0.05, **p<0.005, ***p<0.0005 compared to controls using a two-way ANOVA with Tukey’s correction. (C) Representative images of SEP::GLR-1 fluorescence in control and *mcu-1(lf)* animals at AVA nerve ring for pre-training and 4-hours post-training time points. (D) Total fluorescence (a.u.) or (E) maximum fluorescence of SEP::GLR-1 at AVA nerve ring in control (n ≥ 15) and *mcu-1(lf)* (n ≥ 14) animals for pre-training and 4-hours post-training time points. Data is represented as mean ± sem; ns, not significant, *p<0.05 compared to controls using a one-way ANOVA with Dunnett’s test.

To understand when during training mitochondrial calcium uptake is necessary for plasticity and behavioral sensitization, we pharmacologically blocked MCU-1-dependent calcium influx using ru360 (Figure 3B),^85^ a selective MCU-1 inhibitor we have previously used in *C. elegans.*^83^ Acute ru360 treatment immediately before optogenetic training eliminated reversal sensitization 4-hours after training (Figure 3B). On the other hand, ru360 treatment directly after training had no effect on reversal sensitization (before training ru360: 0.378 ± 0.037, after training ru360: 0.600 ± 0.048 rev/stim). Thus, although behavioral sensitization occurs 4-hours after optogenetic training, activity-dependent MCU-1 calcium uptake is necessary during the training period.

We have previously shown that activity-dependent mitochondrial calcium uptake via MCU-1 upregulates mitoROS signaling which regulates glutamate transport and synaptic insertion,^83^ so we next investigated whether postsynaptic MCU-1 is necessary for changes in synaptic GLR-1 observed following optogenetic training (Figure 2). To do this, we expressed both AVA SEP::GLR-1 and ASH ChR2 in the wildtype or *mcu-1(lf)* background (Figure 3C). Following optogenetic training and subsequent sensitization, we again observed a significant increase in total synaptic GLR-1 abundance (Pre: 210.4 µM^2^ ± 27.43, 4Hr PT: 583.1 ± 118.4; Figure 3D) and maximal intensity (Pre: 273.5 ± 27.07 to 4Hr PT: 673.4 ± 105.8; Figure 3E) in wildtype controls. In contrast, *mcu-1(lf)* animals did not exhibit any change in synaptic GLR-1 abundance or maximum intensity (Figures 3C-E), aligning with the lack of sensitization after training as seen in reversal behavior (Figure 3A-B). Together, these results show that MCU-1 function in AVA is necessary for reversal sensitization and recruitment of GLR-1 to presumptive ASH-AVA synapses is associated with reversal sensitization 4-hours post-training.

### Postsynaptic GLR-1 activation drives mitoROS increase in synapse-adjacent mitochondria after optogenetic training

Neuronal mitochondria adapt to handle activity-driven functions through modulation of morphology, by fusion and fission,^12,17,22^ and by altering ATP output and subsequent ROS production.^4,5^ Mitochondrial calcium uptake via MCU-1 directly contributes to these essential functions by driving morphology changes and energy adaptations.^86^ If and how these adaptations contribute to postsynaptic changes that lead to synaptic plasticity have not been well studied. After discovering that postsynaptic MCU-1 function is necessary for reversal sensitization, we were interested in analyzing mitochondrial morphological and functional dynamics throughout the postsynaptic neuron during sensitization. For this analysis, we quantified mitochondrial shape and local ROS fluctuations using the genetically encoded ROS sensor, roGFP.^87^ This ratiometric sensor utilizes a reversible ROS-sensitive fluorescence change at 405nm (measuring oxidation) and 488nm (used to normalize expression).^88,89^ As described previously,^83^ we targeted roGFP to the outer mitochondrial membrane (mito-roGFP) to analyze mitochondria morphology dynamics and quantify local mitoROS levels across the AVA neuron before and after training (Figure 4A). Areas imaged include the soma, nerve ring (direct ASH input), bifurcation (no ASH synapses present), and below the bifurcation (indirect ASH input; Figure 4B). Interestingly, we observed a post-training mitoROS spike at somatic (405nm/488nm ratio: 0.188 ± 0.047 to 0.318 ± 0.038) and nerve ring mitochondria (405nm/488nm ratio: 0.053 ± 0.010 to 0.104 ± 0.025; Figures 4C-F). In addition, postsynaptic mitoROS increases dissipated by 4-hours post-training in wildtype animals, consistent with the requirement for MCU-1 function during, but not after, training (Figure 3B). This increase in mitoROS was not seen in *mcu-1(lf)* animals (Figures 4C-F). Interestingly, mitochondria at synapses near the AVA bifurcation showed a decrease after training and 4-hours post-training (Figure S4A-B) and mitochondria below the AVA bifurcation showed a slight increase at 4-hours post-training (Figure S4C-D). Thus, mitochondrial proximity to ASH inputs is an important determinant in the change and timing of mitoROS production.

**Figure 4:**
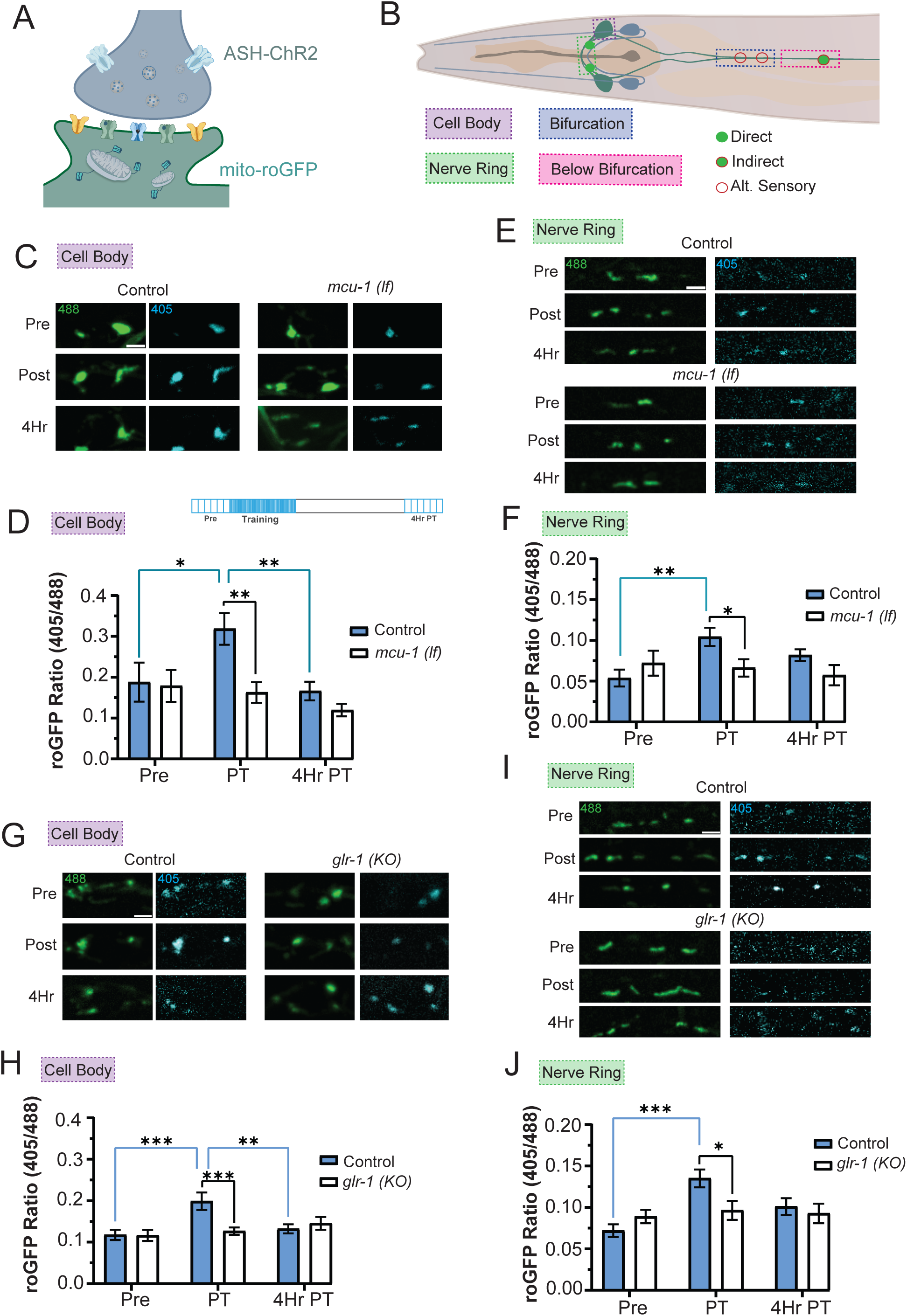
Repetitive neuronal activation with optogenetic training increases mitochondrial ROS production in an MCU-1 dependent manner. (A) Illustration of presynaptic ChR2 and postsynaptic environment with roGFP-tagged mitochondria (mito-roGFP). (B) Illustration displaying AVA interneuron imaging locations compared to ASH sensory neuron. (C and E) Representative images of mito-roGFP fluorescence at AVA (C) cell bodies or (E) nerve ring in a single Z-plane when excited with 488 nm or 405 nm light at pre-training, post-training, and 4-hours post-training time points. (D and F) Mito-roGFP fluorescence ratio (405/488 nm) at AVA (D) cell bodies in control (n = 9 cell bodies from 9 animals) and *mcu-1(lf)* (n = 7 cell bodies from 7 animals) animals at pre-training, post-training, and 4-hours post-training time points. (F) Mito-roGFP fluorescence ratio (405/488 nm) at AVA nerve ring in control (n ≥ 12 mitochondria from 6 animals) and *mcu-1(lf)* (n ≥ 17 mitochondria from 7 animals) animals for pre-training, post-training, and 4-hours post-training time points. (G and I) Representative images of mito-roGFP fluorescence at AVA (G) cell bodies or (I) nerve ring in a single Z-plane when excited with 488 nm or 405 nm light at pre-training, post-training, and 4-hours post-training time points. (H) Mito-roGFP fluorescence ratio (405/488 nm) at AVA cell bodies in control (n ≥ 14 cell bodies from ≥ 7 animals) and *glr-1(KO)* (n ≥ 10 cell bodies from ≥ 9 animals) animals for pre-training, post-training, and 4-hours post-training time points. (J) Mito-roGFP fluorescence ratio (405/488 nm) at AVA nerve ring in control (n ≥ 14 mitochondria from ≥ 7 animals) and *glr-1(KO)* (n ≥ 26 mitochondria from ≥ 9 animals) animals for pre-training, post-training, and 4-hours post-training time points. Scale bars = 2 µm. Data is represented as mean ± sem; ns: not significant, *p<0.05, **p<0.005, ***p<0.0005 compared to controls using a one-way ANOVA with Dunnett’s test.

As previously mentioned, changes in mitochondrial function are often accompanied or caused by morphological changes,^4,5,13^ so we quantified changes in mitochondrial size using the 488nm fluorescence of mito-roGFP from the same mitochondria analyzed for mito-roGFP ratios (in Figure 4C-4F). As described in the methods, we deconvolved mito-roGFP images (using Mean-Shift Super Resolution) to enhance spatial resolution by 20%. In the soma mitochondria form a network rather than distinct single organelles in processes. Thus we quantified the area of this somatic network as the footprint (as was done in Zhang et al.)^90^ and used the size or area of mitochondria in the processes. We found that mitochondria footprint (total area occupied by mitochondria) within the soma was reduced in *mcu-1(lf)* mutants immediately and 4-hours post-training compared to controls (Figure S4E). We also found that there was a trend toward smaller mitochondrial sizes in *mcu-1(lf)* mutants compared to controls, however, mitochondrial size was unaffected by optogenetic training at the nerve ring where activity-dependent changes in mitoROS production were observed (Figure S4E).

To test if GLR-1 expression is necessary for the mitoROS increase, we measured mitoROS levels in the soma and nerve ring of *glr-1(KO)* animals (Figures 4G-J). After optogenetic training, the mito-roGFP fluorescence ratio (405/488 nm) in the soma (Figure 4G-H) and nerve ring (Figure 4I-J) was unchanged in *glr-1(KO)* animals compared to pre-treatment and significantly lower than controls (405/488 nm ratio in soma: 0.117 ± 0.013 to 0.199 ± 0.021, in nerve ring: 0.0719 ± 0.008 to 0.135 ± 0.011).

Thus, GLR-1 is necessary for activity-induced mitoROS signaling in AVA following optogenetic activation of ASH. Then, to determine if *glr-1* and *mcu-*1 are in a linear genetic signaling pathway, we quantified reversal sensitization behavior in a *mcu-1(lf); glr-1(KO)* double knockout (*DKO*). At 4-hours post-training, the *DKO* strain did not show sensitization (*DKO* Pre: 0.233 ± 0.041 vs. 4Hr PT: 0.1444 ± 0.03545; Figure S3E), consistent with both MCU-1 and GLR-1 being necessary for ASH-AVA reversal sensitization. Together, these results indicate that the proximity of mitochondria to active synaptic inputs is associated with increased ROS production and requires cell autonomous MCU-1 function.

### Postsynaptic mitoROS production is necessary and sufficient for reversal sensitization and synapse specific GLR-1 increase

We next wanted to disentangle the effects of MCU-1 dependent mitochondrial calcium influx from effects of mitochondrial ROS signaling,^27,83^ and hypothesized that an increase in mitoROS within the AVA was necessary and sufficient for inducing sensitization. To increase mitoROS levels independent of neurotransmission and calcium influx in a way that would mirror optogenetic training, we targeted the genetically encoded photosensitizer KillerRed (KR) to the outer mitochondrial membrane in AVA (mitoKR; Figure 5A).^65^ Next, we used mito-roGFP in combination with mitoKR to tested different light intensities (data not shown) and activation durations (10 and 60 min, Figure 5A-B) to elicit mitoROS increases that were similar to those following optogenetic activation/training of ASH (see Figures 4D-F), ensuring we could match physiological mitoROS signaling triggered by repeated synaptic activity. Instead of a 1-hour optogenetic activation/training of ASH, we photoactivated mitoKR with orange light (561 nm) for 1-hour. At 4-hours post-training, we activated ASH with blue light to assess reversal sensitization (Figure 5C, Top). Animals were presented with and without AVA mitoKR expression, but all groups expressing ChR2 in ASH, to this new protocol and tested for reversal sensitization as before (Figure 5C). In animals expressing mitoKR, 1-hour of orange light training led to reversal sensitization (0.650 rev/stim ± 0.0520) which was not observed in controls without mitoKR expression (0.440 rev/stim ± 0.0400). These results suggested that mitoROS signaling from postsynaptic mitoKR is sufficient for reversal sensitization. Next, to exclude the possibility that blue light testing during pre-training induces mitochondrial calcium uptake through MCU-1 and, as a result, mitoROS production, we expressed mitoKR in the *mcu-1(lf)* mutant background and performed orange light training. *Mcu-1(lf)* animals with mitoKR showed a significant increase in blue light-induced reversals 4-hours post-training (0.586 rev/stim ± 0.043) compared to *mcu-1(lf)* without mitoKR (0.414 rev/stim ± 0.033; Figure 5C). Taken together, these results demonstrate that mitoROS is necessary and sufficient for reversal sensitization, independent of MCU-1 function. It is important to note that longer photoactivation durations and light intensities for optogenetic mitoKR stimulation (High mitoKR) prevented reversal sensitization and instead led to modest desensitization (Figure 5D). This suggests that positive regulation of reversal behavior by mitoROS signaling may operate within a specific physiological range.

**Figure 5:**
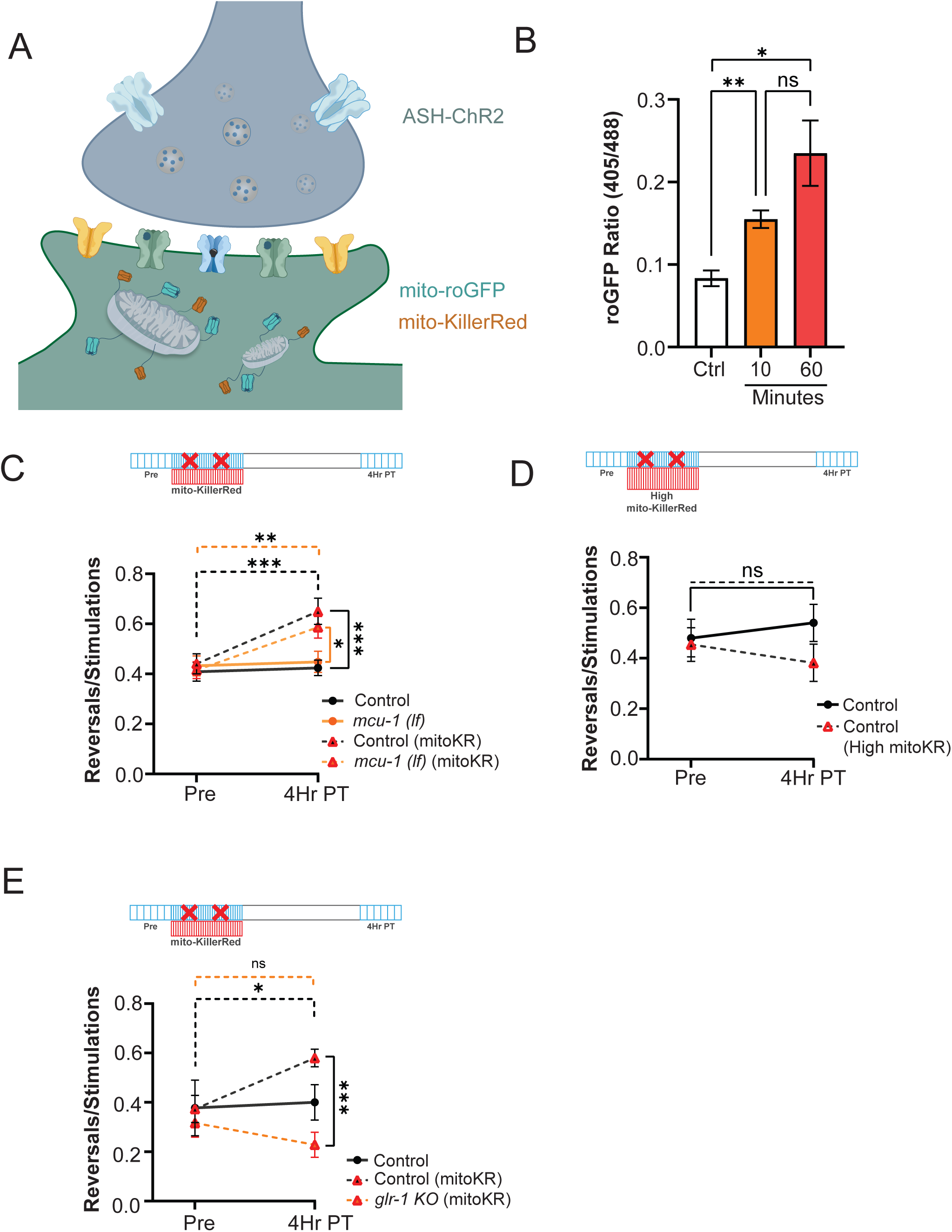
Increased mitochondrial ROS production at postsynaptic sites is necessary and sufficient for reversal sensitization. (A) Illustration displaying pre-synaptic ChR2 and post-synaptic mito-KillerRed and mito-roGFP expression. (B) Mito-roGFP fluorescence ratio (405/488 nm) at AVA nerve ring after 0, 10 or 60 minutes of mitoKR activation (n ≥ 6 mitochondria from 2 animals). Data is represented as mean ± sem; ns: not significant, *p<0.05, ***p<0.0005 compared to controls using a one-way ANOVA with Dunnett’s test. (C) Reversals/optogenetic stimulations ratio in control (n = 24), *mcu-1(lf)* (n = 24), control (mitoKR) (n = 20) and *mcu-1(lf)* (mitoKR) (n = 27) animals for pre-mitoKR and 4-hours post-mitoKR activation time points. (D) Reversals/optogenetic stimulations ratio in control animals for pre-mitoKR (High ROS) and 4-hours post-mitoKR (High ROS) activation time points (n = 11). (E) Reversals/optogenetic stimulations ratio in control (n ≥ 10 animals), control (mitoKR) (n ≥ 9), and *glr-1(KO)* (mitoKR) (n ≥ 12) animals for pre-training and 4-hours post-training time points. Scale bar = 2 µm. Data is represented as mean ± sem; ns: not significant, *p<0.05, **p<0.005, ***p<0.0005 compared to controls using a two-way ANOVA with Tukey’s correction.

Our previous results show that GLR-1 is necessary for reversal sensitization (Figure 1), is increased at nerve ring synapses (Figure 2), and is necessary for mitoROS elevation after optogenetic training (Figure 4). Thus, we hypothesized that GLR-1 is necessary for mitoROS initiation and is a downstream effector of ROS signaling. To test this, we repeated mitoKR training in the *glr-1(KO)* background (Figure 4E) and found that mitoKR activation did not elicit reversal sensitization in *glr-1(KO)* animals. Altogether the data presented in Figure 5 demonstrates that local mitoROS is necessary and sufficient for sensitization after optogenetic training, and that GLR-1 is a key effector of the activity-dependent mitoROS signaling.

## Discussion

In this study, we investigated the role of mitochondrial ROS signaling in the plasticity underlying sensitization of *C. elegans* reversal behavior. In *C. elegans,* reversals are a natural avoidance response to mechanical and chemical stimuli and are largely mediated by direct synaptic transmission from the multimodal ASH neurons to the AVA command interneurons (Figure 1A).^51,67^ The polymodal sensory neurons ASH have been extensively studied for their central role in mediating avoidance responses to noxious chemicals, osmotic pressure and mechanical stimulation.^91–94^ Previous literature has shown that this reversal avoidance behavior involving ASH and AVA is modulated by environmental conditions^70^ through input from other sensory neurons through cGMP and gap-junction connections.^95,96^ Additionally, cross-modal sensitization of ASH has been reported, using reversal length as output and thought to be due to arousal due to an aversive stimulus such as by the release of neuropeptides from touch neurons.^70^ Here, we showed that one hour of optogenetic stimulation of ASH led to sensitization of AVA to presynaptic stimulation 4-hours after training (Figure 1). This avoidance sensitization required transcription and recruitment of ionotropic glutamatergic receptors (GLR-1) to ASH-AVA synapses (Figure 2). Consistent with our prior studies showing activity-dependent mitoROS production, this current study demonstrated that mitoROS signaling is upregulated by optogenetic training and requires postsynaptic MCU-1 function in AVA mitochondria (Figure 3 and 4). Finally, we demonstrated that postsynaptic mitoROS signaling in AVA was necessary and sufficient for increasing the probability of optogenetic-induced reversals (Figure 5). To our knowledge, a direct role for activity-dependent mitoROS signaling in regulating the synaptic plasticity that underlies behavioral sensitization has not been previously described.

Prior studies in hippocampal slices and cell culture have shown that activity-dependent ROS production is important for LTP and other forms of synaptic plasticity.^66^ Specifically, it has been shown that calcium influx through glutamate receptors leads to enhanced ROS generation.^97^ In vertebrate neurons, it is thought that the highly regulated and synapse-localized NAPDH oxidase (NOX) plays a key role in activity-dependent ROS signaling and plasticity.^98,99^ Although there is some evidence in non-neuronal cells that ROS enhances calcium-induced calcium release, evidence in neurons has been lacking.^98^ In *C. elegans* there are no genetic homologues of NOX and no evidence of their presence at synapses, but there are dual oxidases (DUOX1 and DUOX2) found at the plasma membrane of epithelial cells and important for cuticle development and immune responses.^100,101^ Our previous studies have shown that the genetic loss-of-function mutant of the neuronal catalase, *ctl-2,* which increases cytoplasmic ROS, leads to decrease activity-dependent calcium influx and decreases somatic export and delivery of GLR-1 in AVA neurons.^102^ In another study, we showed that acute, local photoactivation of mitoKR (designed to produce ROS at a level comparable to AVA activation for 10 minutes) led to decreased synaptic insertion of GLR-1.^83^ Contrary to these prior results, here we calibrated whole-animal photoactivation of mitoKR to correspond to mitoROS levels measured in somatic and proximal mitochondria after 1-hour of optogenetic training associated with plasticity. In this paradigm mitoROS is a positive signal, necessary and sufficient to induce synaptic strengthening of ASH-AVA synapses and reversal behavior in *C. elegans*.

Many recent studies have now shown that mitochondria play a central role in developmental and adult synaptic plasticity.^6,17,103–105^ A subset of these studies show that cytoplasmic calcium uptake/buffering by mitochondria is central to their role in synaptic plasticity.^18,103^ In this study, we did not measure cytoplasmic calcium but show that mitoROS elevation in soma and proximal synaptic mitochondria depends on MCU-1 function, indirectly implying that calcium signaling is necessary upstream of mitoROS production. However, we show that it can be bypassed by mitoROS, since photoactivation of mitoKR in AVA neurons of *mcu-1(lf)* mutants led to reversal sensitization. In addition, we demonstrate that synaptic increase of GLR-1 necessary for sensitization, is dependent on MCU-1 (Figure 3C). Thus, although mitoROS from photoactivation seems sufficient for reversal sensitization, it requires activity-dependent calcium signaling in physiological conditions. Our data shows that transcription is necessary for sensitization, since blocking transcription prevented synaptic increase in GLR-1. However, this does not exclude that other transcripts may also play a role in this paradigm. Since mitoROS signaling was sufficient to induce reversal sensitization, this implies that ROS signaling is acting on transcription. However, we have not identified which transcription factors and how this signaling leads to GLR-1 transcription elevation.

In summary, our study shows a new role for cell autonomous, activity-dependent mitoROS signaling in regulating synaptic plasticity of avoidance behavior sensitization in *C. elegans*. Our results are limited to the analysis of *C. elegans* avoidance behavior and the specific neurons (ASH-AVA) studied. In addition, both the training and testing of the behavior relied on optogenetic activation of ASH. Nevertheless, prior studies have shown that optogenetic activation of this polymodal nociceptive neuron corresponded to physiological functions of ASH in the avoidance circuit, as well as reflected the role of GLR-1 and glutamatergic transmission in this circuit and behavior. Importantly, since mitochondrial ROS production can depend on the metabolic state of tissues and in the whole organism, these results present the intriguing possibility that this plasticity may be modulated by energy or feeding state of the animal and that coupling mitochondrial ROS production to neuronal activity is a general mechanism for modulating synaptic activity.

## Acknowledgments

We thank Sasha de Henau for the mito-roGFP plasmid, Attila Stetak for the GCaMP6f plasmid, Peter Juo for the ASH-ChannelRhodopsin2 strain. *C. elegans* strains were purchased from the Caenorhabditis Genetics Center, which is funded by the NIH Office of Research Infrastructure Programs (P40 OD010440). This work was largely supported by an R01 awarded to FJ Hoerndli from NIH/NINDS (NS115947).

## Methods

### Plasmid construction

See table Appendix 1 - key resources table for details regarding plasmids used throughout this study. Plasmids were created using In-Fusion Cloning (Takara Bio). DNA primers were created using In-Fusion Primer Design (Takara Bio) and SnapGene.

C. elegans strains

See table Appendix 1 – key resources table for details regarding strains used throughout this study. *C. elegans* strains were maintained under standard conditions (NGM with OP50, at 20℃).^106^ All animals used were 1-day-old adult hermaphrodites selected 24-hours prior to experiments at the L4 stage. Transgenic strains were created by microinjection^107^ including multi copy arrays and single copy insertions using rRMCE.^84^ All strains used in optogenetic experiments were in the *lite-1* null background (allele: ce314 or ok530) to limit light-induced activation of other neurons.^108^

### Confocal microscopy

Imaging was performed using a Yokogawa CSUX1 spinning disc incorporated into an Olympus IX83 confocal microscope with 405, 488, and 561 nm diode lasers (100-150 mW each; Andor ILE Laser Combiner). Images were captured using an Andor iXon Ultra EMCCD (DU-867) camera and a x100/1.40NA oil objective (Olympus). Devices were controlled remotely for image acquisition using MetaMorph 7.10.1 (Molecular Devices).

### In vivo imaging

One-day-old adult hermaphrodites were mounted for confocal imaging by placing a single worm on an agar pad (10% agarose dissolved in M9 buffer) on a microscope slide with 1.8 µl of mounting solution (equal parts polystyrene beadsand 30 mM muscimol). After ∼5 minutes when worm movement slowed, a coverslip was placed onto the pad. The worm’s orientation was adjusted by sliding the coverslip to orient the positioning of the AVA interneurons for imaging.^109^

### Whole-cell neuronal stimulation with Channelrhodopsin 2

Worms from strains expressing Channelrhodopsin 2 were passaged at the L4 stage onto an NGM/OP50 plate coated with 100 µM of all-trans-Retinal (diluted with M9 buffer) and left overnight. Stimulation of strains expressing ChR2 was conducted using an LED array (470 nm, CoolBase 7 LED module LuxeonStar) mounted in a WormLab imaging apparatus. Plates were stimulated 1 second every minute at 200 µW/mm^2^ (adjusted using a custom potentiometer in combination with a digital optical power console (ThorLabs, PM100C) and photodiode sensor (ThorLabs, S170C)) for 60 minutes during optogenetic training and 5 minutes during imaging streams.

### Reversal behavior analysis

WormLab image streams were taken pre-training, post-training, 1-hour post-training, and 4-hours post-training using five activations (1 per minute using the same methodology as training above). Reversals of individual animals were scored at each stimulation (potential activation) over the course of the stream. Reversals were counted if the animal was moving forward or stopped during the onset of the stimulation and proceeded to move in the reverse direction within 2 seconds of activation. Individual animal reversal scores were divided by total stimulations to obtain a reversals/stimulations ratio.

### MitoGCaMP imaging and analysis

The AVA bifurcation was located based on the features described above and monitored to compare postsynaptic *mcu-1* rescue constructs. Fluorescent image streams were collected with a 488 nm imaging laser (power 0.1%; attenuation 10) and taken across the entirety of the AVA neuron (100 ms exposures). Max fluorescence and total activity were analyzed for each mitochondrion as described previously.^83^ Total activity is a combined measure of the amplitude and duration of calcium flux.

### Ratiometric fluorescence imaging and analysis of mito-roGFP

Imaging roGFP occurred at the same time points that optogenetic behavior streams were taken. At each timepoint, worms were removed from the training plate and mounted for imaging within 30 minutes following the *in vivo* imaging protocol above. Animals were first positioned for imaging the AVA cell body, nerve ring, bifurcation, or below bifurcation. After locating mitochondria, image Z-stacks were collected with a 500 ms exposure (100 ms for cell body images) in 0.25 µm steps to capture the entire soma or neuronal process. The 525 nm emission was imaged with 405 nm, then 488 nm illumination at each Z-plane. Max fluorescence from 405 and 488 nm excitation was measured at individual mitochondrion using region measurements (MetaMorph) in a single Z-plane where the 488 nm excitation was highest. The background fluorescence near each mitochondrion was also collected for both excitations. Following logged max intensity values and background subtraction, 405nm/488nm ratios were calculated.

### Mitochondria morphology analysis

Raw 488nm images from mito-roGFP outputs were processed in FIJI using Mean-Shift Super Resolution (MSSR) to the extend spatial resolution of each image followed by Squassh segmentation.^110,111^ Most importantly, this allowed for more complete separation of closely neighboring mitochondria. 15-frame-stacks were taken from raw image stacks containing the mitochondria. Background was subtracted using a rolling ball algorithm and images were then run through one order of MSSR analysis. Output images were then run through the Squassh plugin and manually checked to confirm proper segmentation of mitochondria. To measure mitochondria networks residing in the soma, we used the mitoMAPR plugin to analyze footprints/networks (area) instead of individual mitochondria.^90^

### GLR-1 imaging and analysis

Imaging SEP occurred at the same time points that optogenetic behavior streams were taking, akin to roGFP imaging above. Strains expressing SEP::GLR-1 were removed from the training plate and mounted for nerve ring imaging within 30 minutes following the in vivo protocol above. An image Z-stack of the entire AVA nerve ring was captured in 0.25 µm steps using the 488 nm excitation laser. Fluorescence intensities of SEP puncta were measured using a linescan measurement in MetaMorph followed by a measurement of background intensity. Linescans were then analyzed with a custom MATLAB script.^58^

### MitoKillerRed activation

Stimulation of strains expressing mito-KillerRed (mitoKR) was conducted using an LED array (561 nm, CoolBase 7 LED module LuxeonStar) mounted in a WormLab imaging apparatus. Plates were stimulated for 10 second every minute at 142 µW/mm^2^ (adjusted using a custom potentiometer in combination with a digital optical power console (ThorLabs, PM100C) and photodiode sensor (ThorLabs, S170C) for 60 minutes during optogenetic training. High ROS optogenetic training (Figure 5D) consisted of a 20-second stimulation every minute at 214 µW/mm^2^ for 60 minutes.

### Pharmacological inhibition of MCU-1 with ru360

Ru360 (Sigma-Aldrich, Cat #557440) was reconstituted in water at a concentration of 2 mM, then distributed into 15 µL aliquots (in light safe microcentrifuge tubes) and stored at 4 ℃. Before treatment, an aliquot was diluted to 100 µM with M9 buffer. Animals were then placed on an NGM plate with OP50 and 200 µL of 100 µM ru360 solution was pipetted over the animals, covering the lawn. Treatment was applied for 10 minutes, after which animals were either optogenetically trained or video recorded.

### Pharmacological inhibition of transcription with Actinomycin D

Actinomycin D concentrations and treatment times (Figure S2A) were tested using hsp-16.2::eGFP (ST66, ncIs17).^80^ Plates containing the animals were placed at 37°C for 20 minutes. 40 minutes after heat shock, pharynx fluorescence intensity was imaged and compared to control animals. Actinomycin D (Sigma-Aldrich, Cat #50-76-0) was reconstituted in DMSO at a concentration of 20 mg/mL 5 µL of 20 mg/mL Actinomycin D was then added to 95 µL M9 buffer to make a 0.797 mM solution, stored in dark Eppendorf tubes for light sensitivity. Animals on NGM plates with OP50 were treated with 100 µL of 0.797 mM (1000 μg/mL) Actinomycin D solution by pipetting the solution over the animals, covering the OP50 lawn. After drying, plates were then used in optogenetic training paradigm for behavior and imaging.

### Quantification and statistical analysis

Data are represented as mean ± standard error of the mean (SEM). Specific p-value calculations are listed in the figure legend for each data set. For manual quantification of behavior (reversal counting), data sets were blinded to genotypes and experimental conditions. Outliers were removed from datasets using the ROUT method (Q = 1%).

### Image and data presentation

Graphical illustrations and significance were obtained with GraphPad Prism (Version 10), exported as an enhanced metafile and compiled using Adobe Illustrator (2025). Representative images were selected based on averages from each data set. Postprocessing was used after analysis to better visualize quantifications. Image processing was done using FIJI. All images in each panel were processed identically at the same time.

