## Supplemental figures and information for "Activity-Dependent Postsynaptic Mitochondrial ROS Signaling Drives Avoidance Plasticity *in C. elegans*"

A

ASH/AVA GCaMP
Example Video with
calcium trace

B

Cropped single
reversal video

C

Reversal Example Videos
(sped up) (Pre/4HrPT)

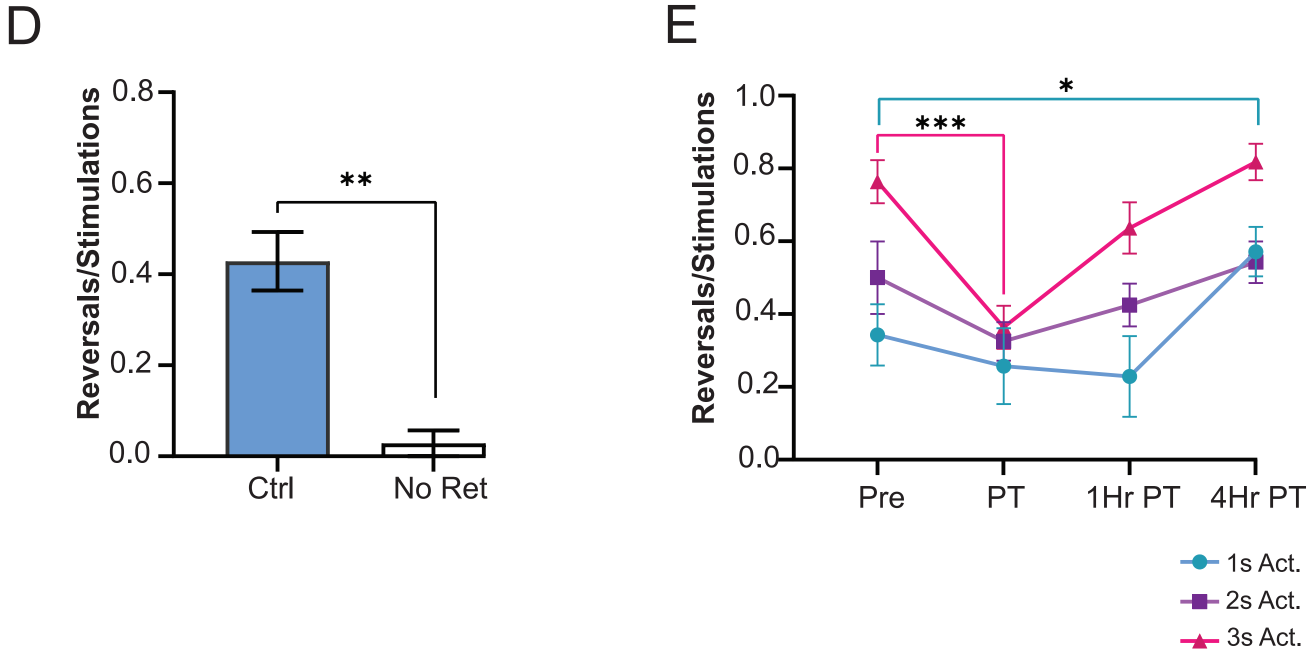

**Supp. Figure 1**
(A) Video showing changes in AVA cytoplasmic GCaMP fluorescence following optical activation of ASH::ChR2 on a confocal microscope (top) and a representative GCamp trace (bottom). (B) Video displaying an example reversal following optogenetic activation of ASH::ChR2 on a plate. (C) Videos showcasing reversals pre-training and 4-hours post-training with 470nm light activations.These representations are displayed at 3x speed with activations at 10/30/50/70/90 seconds (original videos have initial activation start at 30-sec and repeat every minute over 5-min). (D) Reversals/stimulations ratio in controls and no retinol animals (n = 7 animals). Data is represented as mean ± sem; ns, not significant, $*p<0.05$ compared to controls using an unpaired Mann-Whitney test. (E) Reversals/stimulations for control animals using 1, 2, or 3 second activation times ($n \geq 7$). Data is represented as mean ± sem; ns, not significant, $*p<0.05$, $**p<0.005$ compared to controls using a one-way ANOVA with Tukey's correction.

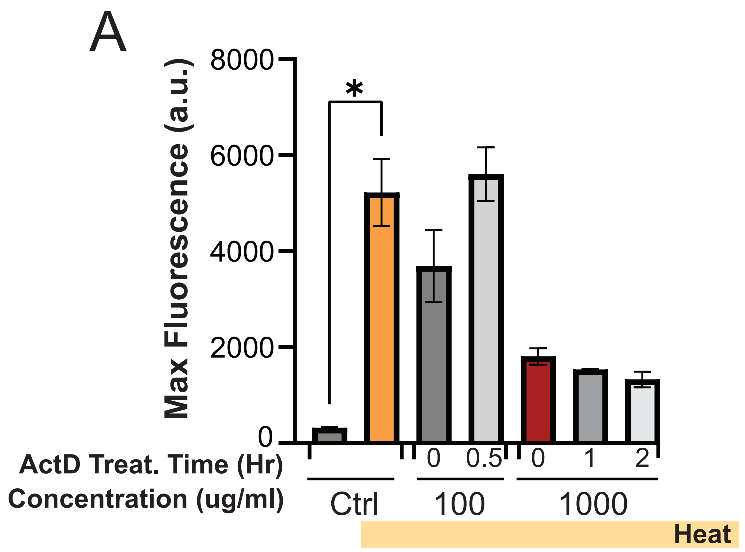
A
Max Fluorescence (a.u.)
8000
6000
4000
2000
0
*
ActD Treat. Time (Hr)
0
0.5
0
1
2
Concentration (ug/ml)
Ctrl
100
1000
Heat

**Supp. Figure 2**

(A) GFP fluorescence intensity in pharynx muscle from hsp-16.2::eGFP animals after actinomycin D treatments of various concentrations and treatment durations with and without heat shock as indicated ($n \geq 2$ animals). Data is represented as mean ± sem; ns: not significant, *$p<0.05$ compared to controls using a one-way ANOVA with Dunnett's test.

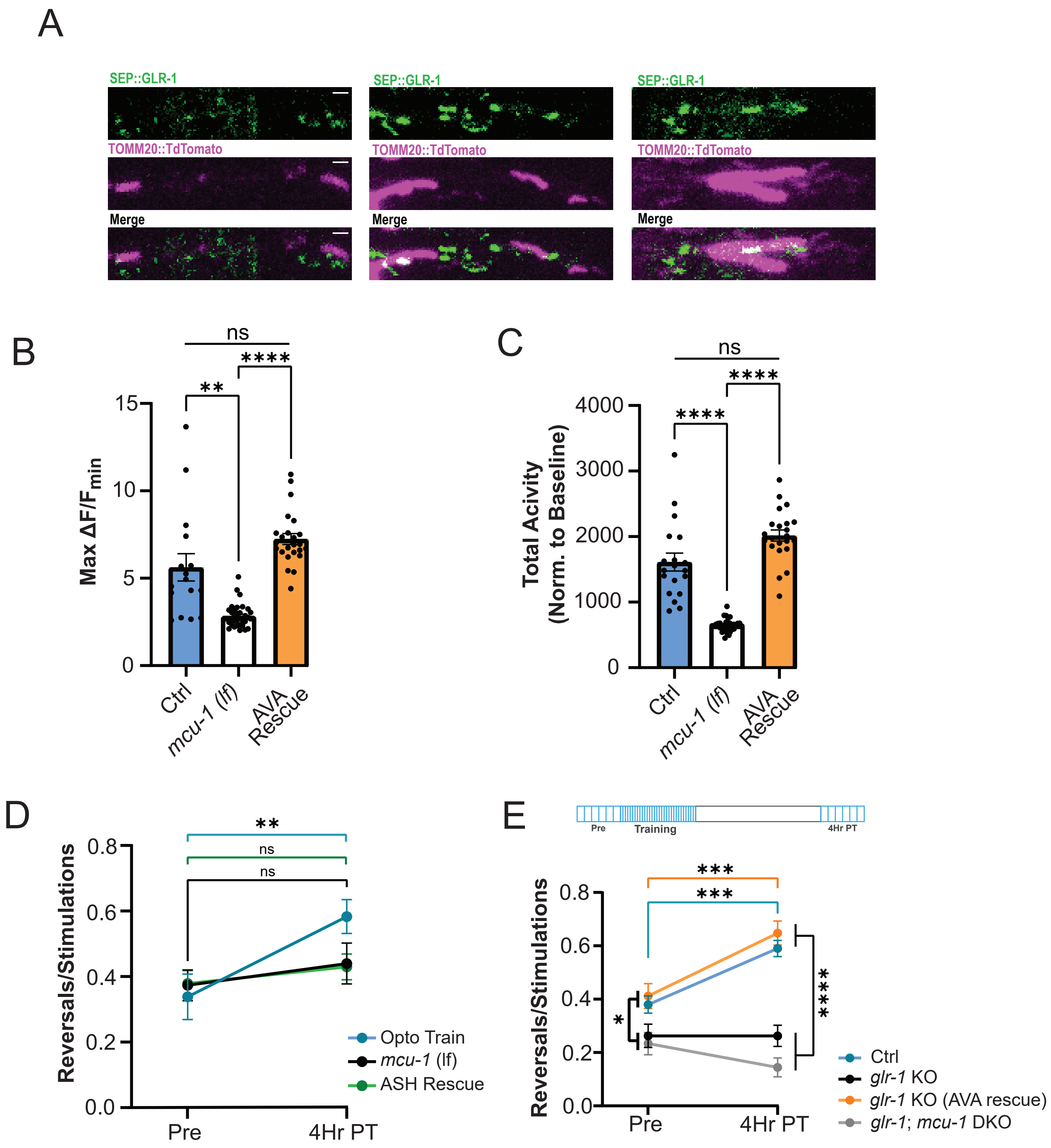
A
SEP::GLR-1
TOMM20::TdTomato
Merge
B
ns
**
****
Max ΔF/F min
Ctrl
mcu-1 (lf)
AVA Rescue
C
ns
****
****
Total Acivity (Norm. to Baseline)
Ctrl
mcu-1 (lf)
AVA Rescue
D
**
ns
ns
Reversals/Stimulations
Pre
4Hr PT
Opto Train
mcu-1 (lf)
ASH Rescue
E
Pre
Training
4Hr PT
***
***
*
****
Reversals/Stimulations
Ctrl
glr-1 KO
glr-1 KO (AVA rescue)
glr-1; mcu-1 DKO

**Supp. Figure 3**

(A) Representative images of tdTomato::TOMM-20 (mitomarker) and SEP::GLR-1 fluorescence in AVA nerve ring of control animals. Scale bar = 2 µm (B-C) Basal mitoGCaMP calcium measurements including (A) MaxF/Fmin and (B) Total Activity/Baseline (n ≥ 16 mitos in ≥4 animals). Data is represented as mean ± sem; ns: not significant, **p<0.005, ***p<0.0005, ****p<0.00005 compared to controls using a one-way ANOVA with Dunnett's test. (D) Reversals/optogenetic stimulations ratio in controls (n ≥ 12), mcu-1*(lf)* (n = 15), and *mcu-1(lf)* (ASH Rescue) (n ≥ 27) animals at pre-training and 4-hours post-training time points. (E) Reversals/optogenetic stimulations ratio in control (n ≥ 15), *glr-1(KO)* (n = 16), *glr-1(KO)* (AVA Rescue) (n ≥ 17), and *mcu-1(lf)*; *glr-1(KO)* (n ≥ 12) animals at pre-training and 4-hours post-training time points. Data is represented as mean ± sem; ns: not significant, **p<0.005, ***p<0.0005, ****p<0.00005 compared to controls using a two-way ANOVA with Tukey's correction.

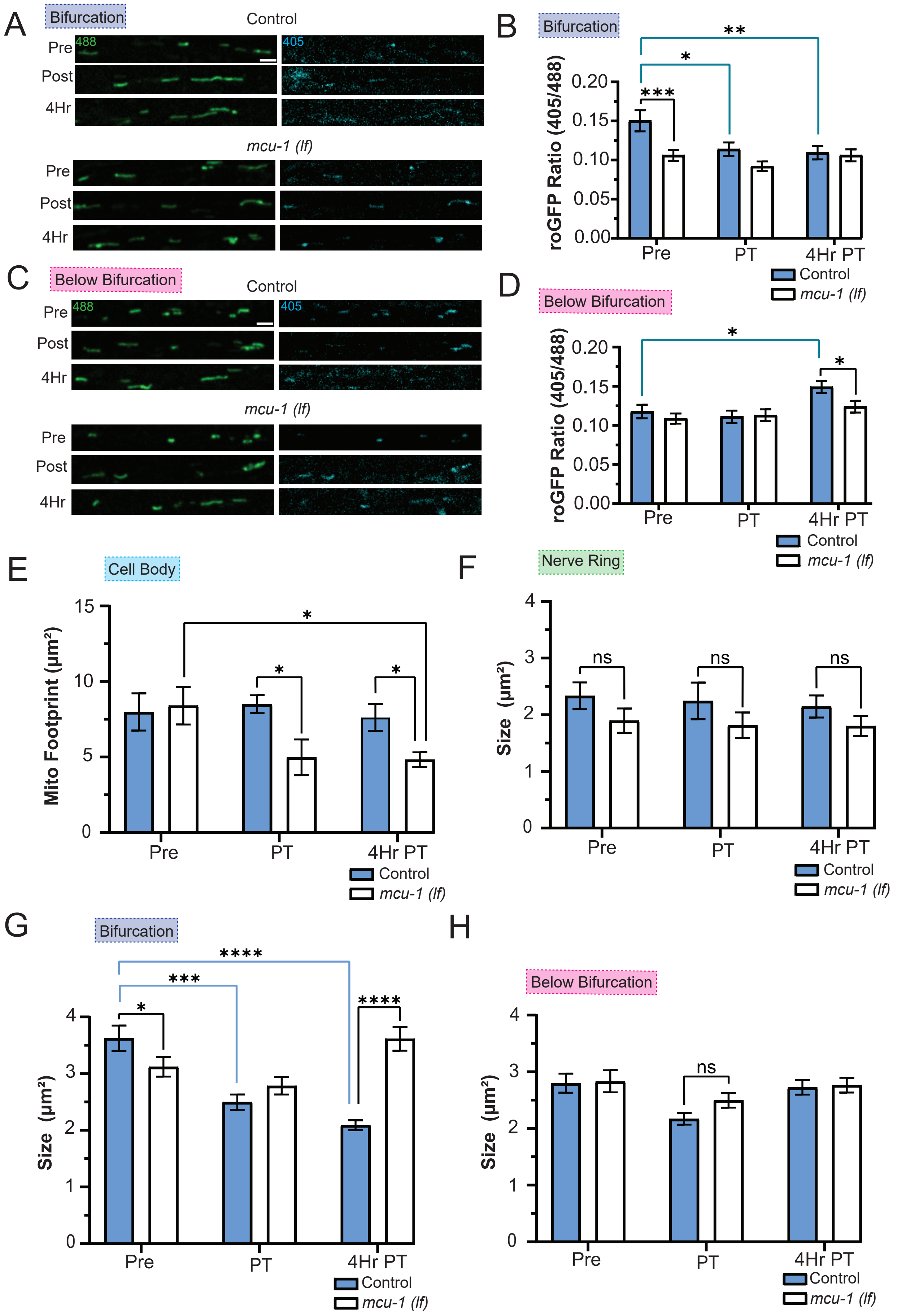

A
Bifurcation
Control
Pre
Post
4Hr
488
405
mcu-1 (lf)
B
Bifurcation
roGFP Ratio (405/488)
0.20
0.15
0.10
0.05
0.00
***
*
**
Pre
PT
4Hr PT
Control
mcu-1 (lf)
C
Below Bifurcation
D
Below Bifurcation
E
Cell Body
Mito Footprint (μm²)
15
10
5
0
F
Nerve Ring
Size (μm²)
ns
G
Bifurcation
****
H
Below Bifurcation

**Supp. Figure 4**

(A and C) Representative images of mito-roGFP fluorescence (A) at or (C) below AVA bifurcation in a single Z-plane when excited with 488 nm or 405 nm light at pre-training, post-training, and 4-hours post-training time points. Scale bar = 2μm. (B) Mito-roGFP fluorescence ratio (405/488 nm) at AVA bifurcation in control (n ≥ 52 mitochondria from 6 animals) and *mcu-1(lf)* (n ≥ 64 mitochondria from 7 animals) animals for pre-training, post-training, and 4-hours post-training time points. (D) Mito-roGFP fluorescence ratio (405/488 nm) below AVA bifurcation in control (n ≥ 56 mitochondria from 6 animals) and *mcu-1(lf)* (n ≥ 72 mitochondria from 7 animals) animals at pre-training, post-training, and 4-hours post-training time points. (E) Mito footprint ($\mu m^2$) at AVA cell bodies in control (n = 9 cell bodies from 9 animals) and *mcu-1(lf)* (n = 7 cell bodies from 7 animals) animals for pre-training, post-training, and 4-hours post-training time points. (F-H) Mito size ($\mu m^2$) at AVA nerve ring in control (n ≥ 12 mitochondria from 6 animals) and *mcu-1(lf)* (n ≥ 17 mitochondria from 7 animals) animals for pre-training, post-training, and 4-hours post-training time points. (G) Mito size ($\mu m^2$) at AVA bifurcation in control (n ≥ 52 mitochondria from 6 animals) and *mcu-1(lf)* (n ≥ 64 mitochondria from 7 animals) animals for pre-training, post-training, and 4-hours post-training time points. (H) Mito size ($\mu m^2$) at AVA below bifurcation in control (n ≥ 56 mitochondria from 6 animals) and *mcu-1(lf)* (n ≥ 72 mitochondria from 7 animals) animals for pre-training, post-training, and 4-hours post-training time points. Data is represented as mean ± sem; ns: not significant, *p<0.05, **p<0.005, ***p<0.0005, ****p<0.00005 compared to controls using a one-way ANOVA with Dunnett's test.
